# Consensus scHPF Identifies Cell Type-Specific Drug Responses in Glioma by Integrating Large-Scale scRNA-seq

**DOI:** 10.1101/2023.12.05.570193

**Authors:** Hanna Mendes Levitin, Wenting Zhao, Jeffrey N. Bruce, Peter Canoll, Peter A. Sims

## Abstract

Single-cell transcriptomic analyses now frequently involve elaborate study designs including samples from multiple individuals, experimental conditions, perturbations, and batches from complex tissues. Dimensionality reduction is required to facilitate integration, interpretation, and statistical analysis. However, these datasets often include subtly different cellular subpopulations or state transitions, which are poorly described by clustering. We previously reported a Bayesian matrix factorization algorithm called single-cell hierarchical Poisson factorization (scHPF) that identifies gene co-expression patterns directly from single-cell RNA-seq (scRNA-seq) count matrices while accounting for transcript drop-out and noise. Here, we describe consensus scHPF, which analyzes scHPF models from multiple random initializations to identify the most robust gene signatures and automatically determine the number of factors for a given dataset. Consensus scHPF facilitates integration of complex datasets with highly multi-modal posterior distributions, resulting in factors that can be uniformly analyzed across individuals and conditions. To demonstrate the utility of consensus scHPF, we performed a meta-analysis of a large-scale scRNA-seq dataset from drug-treated, human glioma slice cultures generated from surgical specimens across three major cell types, 19 patients, 10 drug treatment conditions, and 52 samples. In addition to recapitulating previously reported cell type-specific drug responses from smaller studies, consensus scHPF identified disparate effects of the topoisomerase poisons etoposide and topotecan that are highly consistent with the distinct roles and expression patterns of their respective protein targets.

## Introduction

Advances in the scalability of single-cell RNA-seq (scRNA-seq) have motivated the development of numerous computational methods for integrating data across multiple subjects and conditions. Common analytical frameworks include clustering and trajectory or pseudo-temporal inference, often followed by differential expression analysis or the identification of genes that are correlated with pseudo-time^1^. These methods have been augmented by various algorithms for batch correction and modeling inter-subject variability to facilitate integration of datasets originating from complex experimental designs^2,3^, so that cells can be assigned to clusters following harmonization. While these are powerful approaches that have led to numerous discoveries, they can also place problematic structural constraints on the data. For example, while clustering is a facile approach to assigning single-cell profiles to major cellular lineages within a highly diverse population, applying clustering to a more homogenous, individual lineage that is comprised of more subtle subsets or undergoing a cell state transition often results in rather arbitrary segregation.

To address these issues, matrix factorization approaches have been developed to reduce dimensionality and identify gene expression programs that contribute to cellular diversity without imposing such structural constraints. Some of these methods are probabilistic and explicitly model the transcript drop-out and technical noise that are intrinsic to scRNA-seq^4-6^. We reported such a method that adapts the hierarchical Poisson factorization (HPF) algorithm for scRNA-seq applications, which we called single-cell HPF (scHPF)^5^. scHPF decomposes a scRNA-seq count matrix into *K* factors, each of which has a weight for each cell and gene in the dataset. The probabilistic model that underlies scHPF enforces sparsity, leading to highly interpretable factors where the top-ranked genes are often readily identifiable as markers of specific cell types, genes involved in specific biological processes, or nuisance signatures that can be removed prior to downstream analysis.

While scHPF has been used to great effect in numerous studies to identify T cell activation signatures in human tissues^7^, acute responses to drugs^8,9^, and drug resistance mechanisms in cancer^10^, the algorithm has some important shortcomings. Specifically, applying scHPF to datasets involving multiple subjects or conditions can be challenging. Indeed, in one of our early applications to scRNA-seq profiles of resting and activated T cells from multiple tissue sites and organ donors, we constructed scHPF models of each sample individually and identified relationships between the resultant factors rather than integrating the data with a single model^7^. While this approach is effective, it presents challenges for directly comparing gene signatures across individuals or conditions. Another frustrating issue is the requirement that the user pre-select a specific value of *K*. One approach to these problems that has been implemented in other algorithms for non-negative matrix factorization is to construct factor models across a broad range of *K*-values and other parameters and identify factors that recur across these models^11^. In this case, the final value of *K* becomes the number of recurrent factors, which can be refined by a final round of training. In addition to providing a robust approach to determining *K*, the resultant models are more effective for integrating disparate scRNA-seq datasets. Here, we describe a consensus implementation of scHPF and apply it to identify cell type-specific drug responses from patient-derived *ex vivo* models of high-grade gliomas and glioblastoma (GBM) across multiple patients and classes of drugs.

## Results

### Consensus single-cell hierarchical Poisson factorization (scHPF)

scHPF is a generative, Bayesian algorithm for probabilistic factorization of scRNA-seq count matrices that produces highly interpretable factors that indicate gene co-expression signatures^5^. The input to scHPF is an un-normalized, *N x M* count matrix *C* where *N* is the number of cells and *M* is the number of genes. scHPF associates each gene *g* and cell *i* with inverse budgets *η*_*g*_ and *ξ*_*i*_, and models these latent variables with Gamma distributions. In addition, we use a second set of Gamma distributions to model the gene and cell loadings for each factor *k*, which we call *β*_*g,k*_ and *θ*_*i,k*_, respectively, with rates that depend on *η*_*g*_ and *ξ*_*i*_. The expression of each gene in each cell is assumed to follow a Poisson distribution with a rate given by the dot product of the gene and cell loadings across factors. We estimate the posterior distributions of the budgets and factors for a given count matrix using variational inference. Thus, scHPF effectively models the generative process underlying the count matrix as a Gamma-Poisson mixture, which has been shown to be an effective description of the noise in the scRNA-seq data when unique molecular identifiers (UMIs) are applied to mitigate amplification noise and bias^12^. To aid in the interpretation of scHPF factors, we additionally compute gene scores for each gene in each factor and cell scores for each cell in each factor that correct the *β*_*g,k*_ and *θ*_*i,k*_ loadings for coverage differences among genes and cells, respectively. We have shown in numerous studies that ranking genes by their gene scores in each factor greatly aids in the biological interpretation of each factor, while the cell scores enable embedding and visualization of scHPF models^5,7-10^.

Consensus scHPF uses the methodology described above to generate many independent models of a count matrix and clusters the resultant factors to identify robust, recurrent factors from which it learns a single, consensus model (**Figure 1**). This procedure overcomes the highly multi-modal posteriors that arise from complex datasets that integrate multiple experimental runs, subjects, and conditions, and is analogous to consensus methods used in previously reported methods like cNMF^11^ and scCoGAPS^13^. Thus, we can capture highly robust co-expression patterns that appear consistently across scHPF training models with different random initializations, while still preserving the ability to approximate the parameter values that best explain the data.

**Figure 1.**
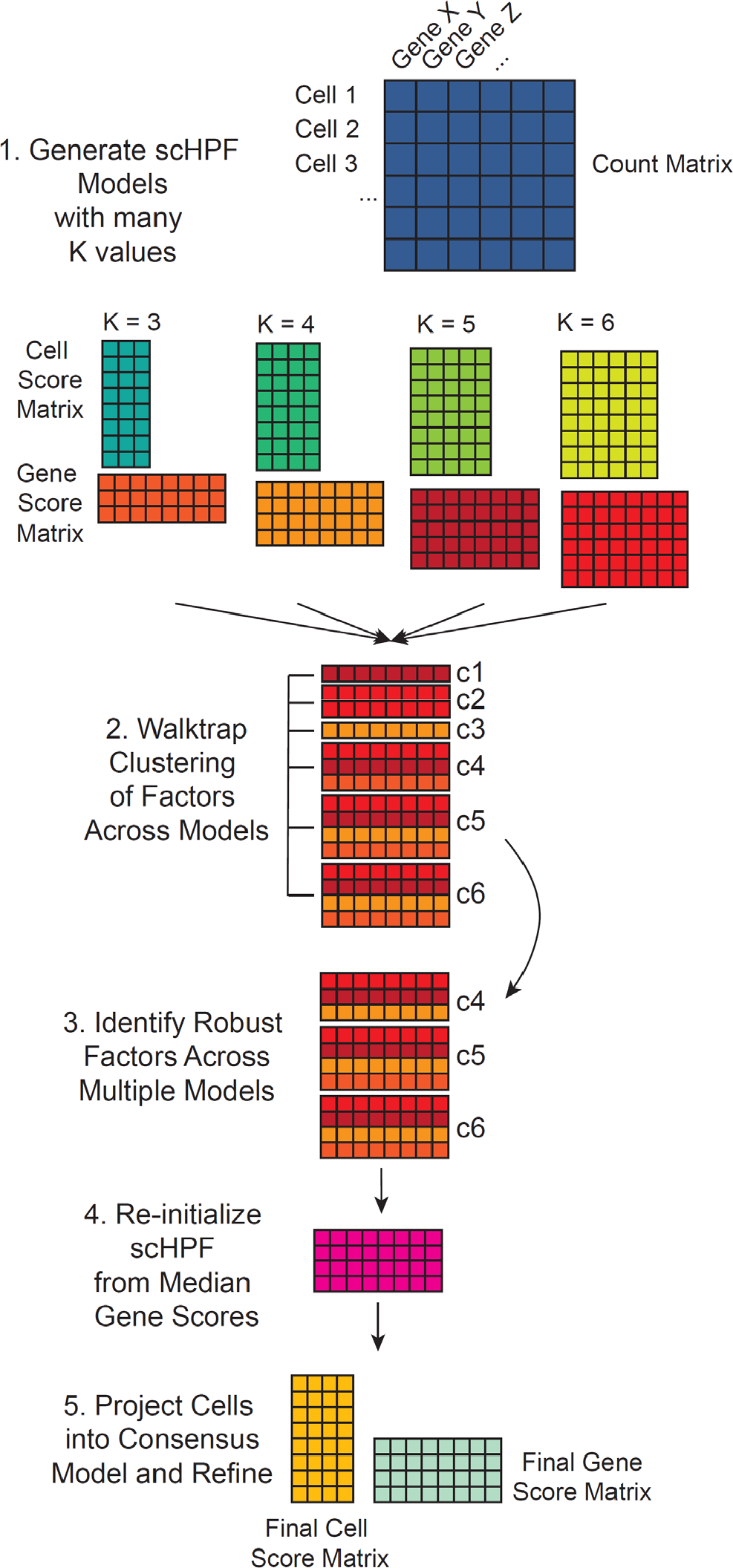
Schematic of consensus scHPF. Starting with an scRNA-seq count matrix, we construct multiple scHPF models with random initializations and different values of *K*. We then cluster the factors across models using their gene score vectors to identify sets of recurrent factors or modules. Finally, we refine a consensus scHPF model by initializing from the median parameters of those modules. The number of modules or recurrent factors becomes the final value of *K*.

After generating scHPF models across different values of *K* and multiple random initializations, we concatenate the gene score matrices across models into a single matrix *X* with factors as columns, genes as rows, and matrix elements as scHPF gene scores. We then identify a subset of genes with highly variable gene scores and cluster the factors using the Walktrap algorithm (see Methods). Clustering allows us to identify factors that recur across models, which gives us the value of *K*. Next, we initialize the variational distributions for the gene weights *β*_*g,k*_ to have shape and rate parameters (*β*_*shape*_ and *β*_*rate*_) equal to the medians of the corresponding variational distributions of each cluster of factors. Similarly, we initialize the variational distributions for *η*_*g*_ to have shape and rate parameters (*η*_*shape*_ and *η*_*rate*_) equal to the median values across the corresponding distributions for all models. Cell-specific variational distributions for *θ*_*i,k*_ and *ξ*_*i*_ are initialized by performing a single round of scHPF updates, while fixing the gene-specific distributions for *β*_*g,k*_ and *η*_*g*_. Finally, we run training updates starting from this initialized consensus model until convergence.

### Identification of Recurrent Drug Responses among Malignantly Transformed Glioma Cells across Diverse Classes of Drugs with Consensus scHPF

Glioblastoma (GBM) is the most common and deadly type of glioma in adults^14^. Standard treatment for GBM at primary diagnosis is surgical resection, but glioma cells infiltrate the brain, inevitably leading to recurrence despite chemotherapy and radiation^14^. GBM heterogeneity is a major obstacle to successful treatment, but detailed studies using scRNA-seq have identified multiple transcriptional states that recur across patients^15,16^. Thus, there is a need for drug screening platforms that can identify cell type-specific drug responses in patient-derived models. We recently reported such a platform that combines drug-perturbed, acute slice culture of GBM surgical specimens with scRNA-seq for identifying cell type-specific drug responses among transformed glioma cells and cells in the tumor microenvironment^8^. We performed detailed studies to credential these *ex vivo* models, demonstrating that they recapitulate the cellular composition and molecular profile of the originating tumor tissue with high fidelity^8^. Here, we combine previously published^8,9^ and newly generated scRNA-seq data from GBM and high-grade glioma slice cultures perturbed with diverse classes of drugs and apply consensus scHPF to identify common and drug-specific gene signatures that occur across 19 patients, 10 treatment conditions, and ∼400,000 individual cells (**Table S1**). To do this, we constructed cell type-specific, consensus scHPF models that integrate the data from all of the patients and drug perturbation conditions in the study for three major cell types – the transformed glioma cells, myeloid cells, and oligodendrocytes. The latter two are non-neoplastic cells in the tumor microenvironment, with myeloid cells comprising the most common infiltrating immune cell type in GBM. This integrated analysis allows us to categorize drugs based on their cell type-specific effects in GBM.

We first performed a global analysis of consensus scHPF models obtained from each of the three major cell types in the dataset. **Figure 2** shows UMAP embeddings of the cell score matrices for each cell type annotated by patient and drug treatment condition. While we expect some degree of clustering by patient in this dataset, because different sets of drug treatments were applied to slice cultures from different patients, there is particularly clear separation by patient for the transformed glioma cells (**Figure 2A,D**). This is consistent with our previous studies and anticipated given the disparate genetic alterations, particularly aneuploidies and other large copy number alterations, that significantly impact gene expression^15^. Similarly, **Figure 3** shows a heatmap of the average cell scores for each scHPF factor in each experimental condition (defined by a patient and a drug perturbation) for the scHPF models for all three major cell types. The columns of the heatmap, which represent patient-drug combinations, are hierarchically clustered, while the rows, which represent factors from the three cell type-specific scHPF models, are grouped by cell type. While the distributions of myeloid cell and oligodendrocyte factors are relatively uniform across patients and drug treatment conditions, there are many scHPF factors that are highly patient- or treatment-specific for the model of transformed glioma cells.

**Figure 2.**
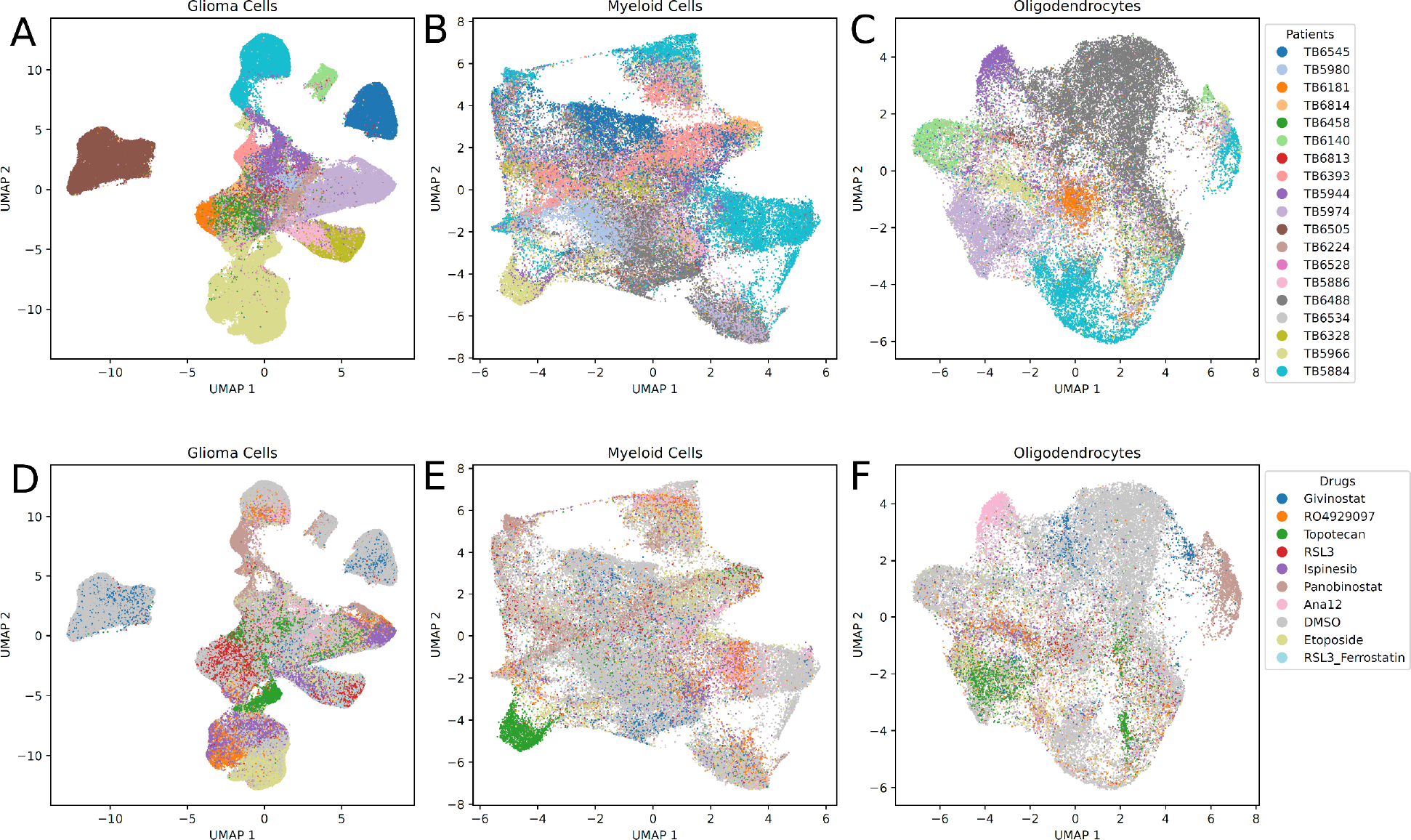
UMAP embeddings of cell score matrices from consensus scHPF models colored by patient ID for A) transformed glioma cells; B) myeloid cells; C) oligodendrocytes; and colored by drug treatment condition for D) transformed glioma cells; E) myeloid cells; F) oligodendrocytes.

**Figure 3.**
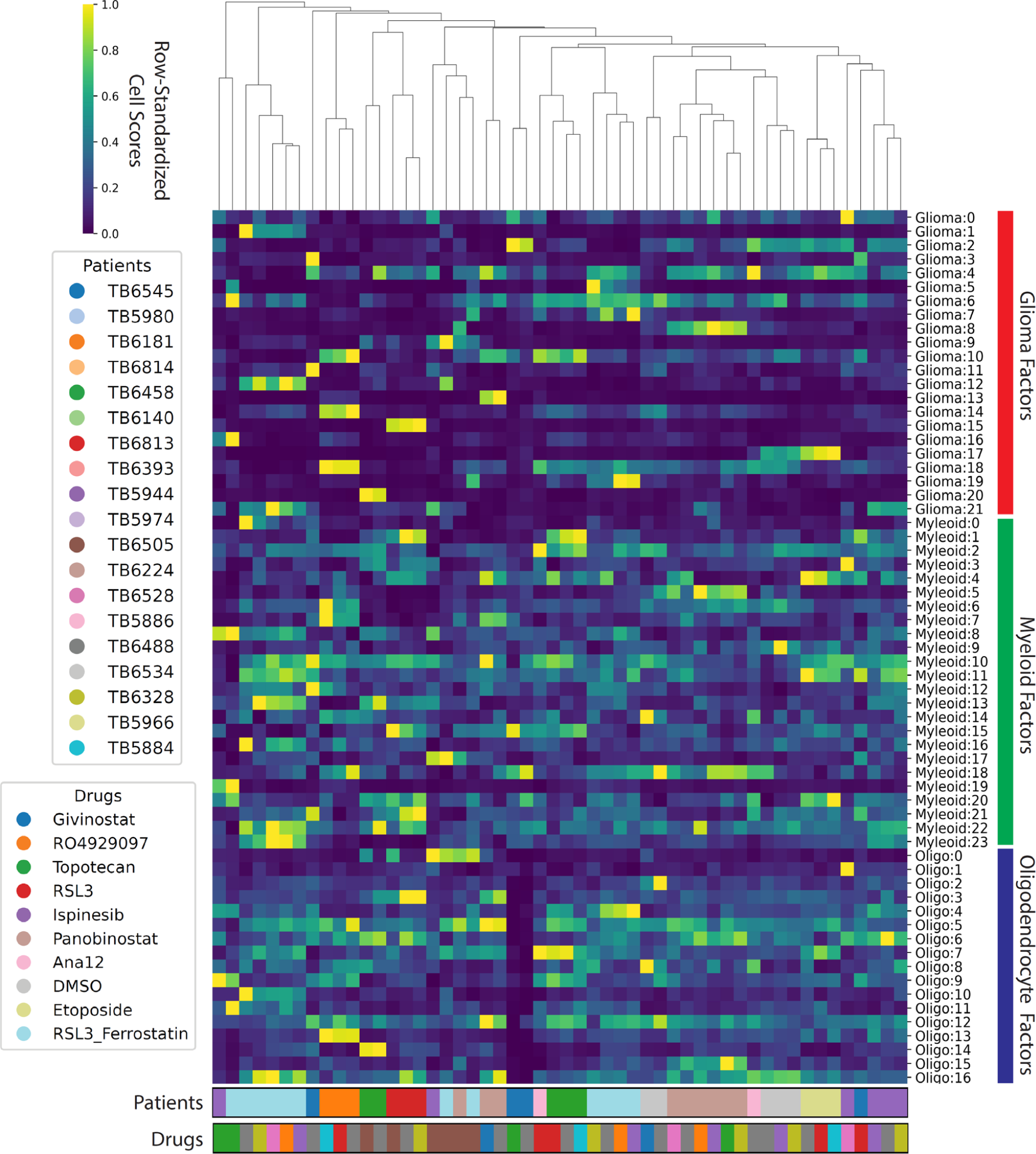
Hierarchically clustered heatmap showing the average cell scores for each patient-drug combination across all factors in the three cell type-specific, consensus scHPF models (glioma cells, myeloid cells, oligodendrocytes).

We next sought to identify scHPF factors with cell scores that were consistently higher or lower across patients for a given drug perturbation relative to the corresponding vehicle controls. To identify these factors, we used the median absolute deviation (MAD) to identify subpopulations with high cell scores for each factor relative to the appropriate control sample (the most adjacent slice culture treated with vehicle control, see Methods). We highlight four particularly interesting signatures in **Figure 4. Figure 4A** shows the treatment effect results across patients and drugs for a factor representing proliferation (*CENPF, TOP2A, MKI67*). As expected for a topoisomerase II poison and consistent with our earlier study^8^, slice cultures from five of the six patients that were treated with etoposide show a decrease in cell frequencies with high cell scores for this factor. Surprisingly, we do not observe this decrease for slice cultures treated with topotecan, which targets topoisomerase I. Also consistent with our previous study^8^, we identify a factor corresponding to metallothionein induction (*MT1G, MT1X, MT1H, MT1E*) that is uniformly elevated in slice cultures treated with the HDAC inhibitor panobinostat (**Figure 4B**). We observe a similar, but slightly attenuated effect, for givinostat, the other HDAC inhibitor in our study.

**Figure 4.**
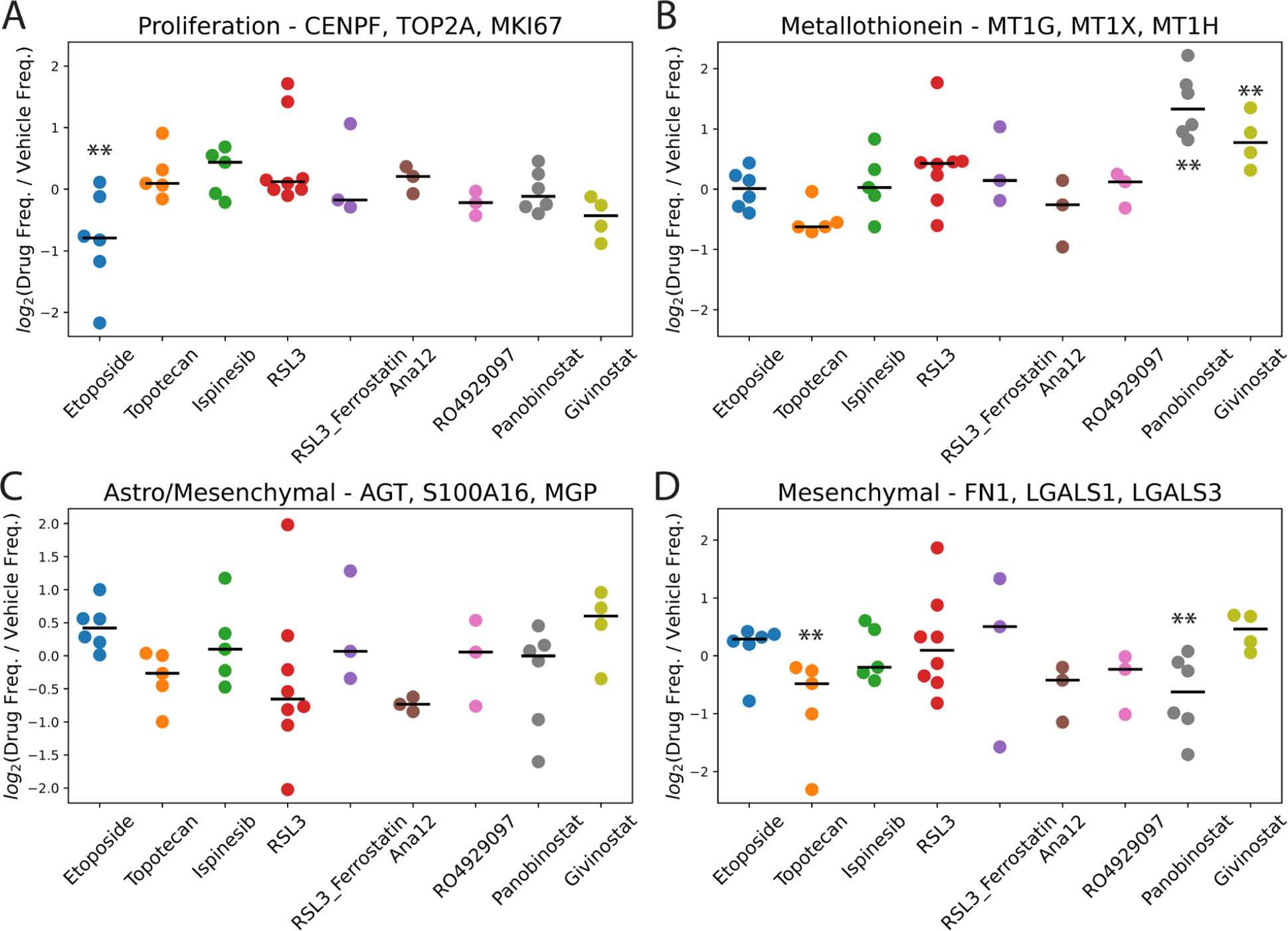
Fold-changes in the frequencies of cells with high cell scores in drug-treated vs. vehicle control-treated slice cultures in the transformed glioma cell scHPF model for A) a proliferation factor; B) a metallothionein factor; C) an astrocyte/mesenchymal factor; and D) a mesenchymal factor. Here, each dot represents an individual patient (i.e. biological replicates). For each drug, ** indicates FDR<0.05 based on a linear mixed model.

A third factor with markers of both astrocyte- and mesenchymal-like glioma cells decreases in most of the slice cultures treated with topotecan, RSL3, and Ana12 (**Figure 4C**). Ana12 targets NTRK2, which is widely expressed across multiple glioma subtypes^17^. While we expected this result for the ferroptosis-inducing GPX4 inhibitor RSL3 based on our prior work^9^, it is somewhat counter-intuitive for topotecan to target a subpopulation that is typically quiescent in glioma. Nonetheless, targeting of astrocyte-like or mesenchymal glioma cells by topotecan and the apparent increase in this factor in the etoposide-treated slices are consistent with their opposite effects on cycling populations shown in **Figure 4A**. Furthermore, we observed an even greater and highly significant impact of topotecan on cell frequencies for a second, highly mesenchymal-like factor (**Figure 4D**).

To further interrogate the disparate effects of the two topoisomerase poisons topotecan and etoposide, we examined the expression patterns of the genes encoding their targets, *TOP1* and *TOP2A*, respectively (**Figure 5**). Interestingly, we observed that while *TOP2A* expression is highly restricted to proliferating glioma cells as evidenced by its co-expression with *MKI67* (**Figure 5D-F**). Conversely, *TOP1* is more pervasively expressed and strongly co-expressed with *CD44*, a marker of mesenchymal glioma cells (**Figure 5A-C**). This finding is consistent with the broader functional role of topoisomerase I, which is critical for both DNA replication during the cell cycle and transcription, regardless of whether or not the cells are cycling^18^. Thus, although etoposide and topotecan are both topoisomerase poisons, we might expect etoposide’s effects to be restricted to proliferating cells, while topotecan is able to target broader populations that include more quiescent astrocyte-like and mesenchymal glioma cells.

**Figure 4.**
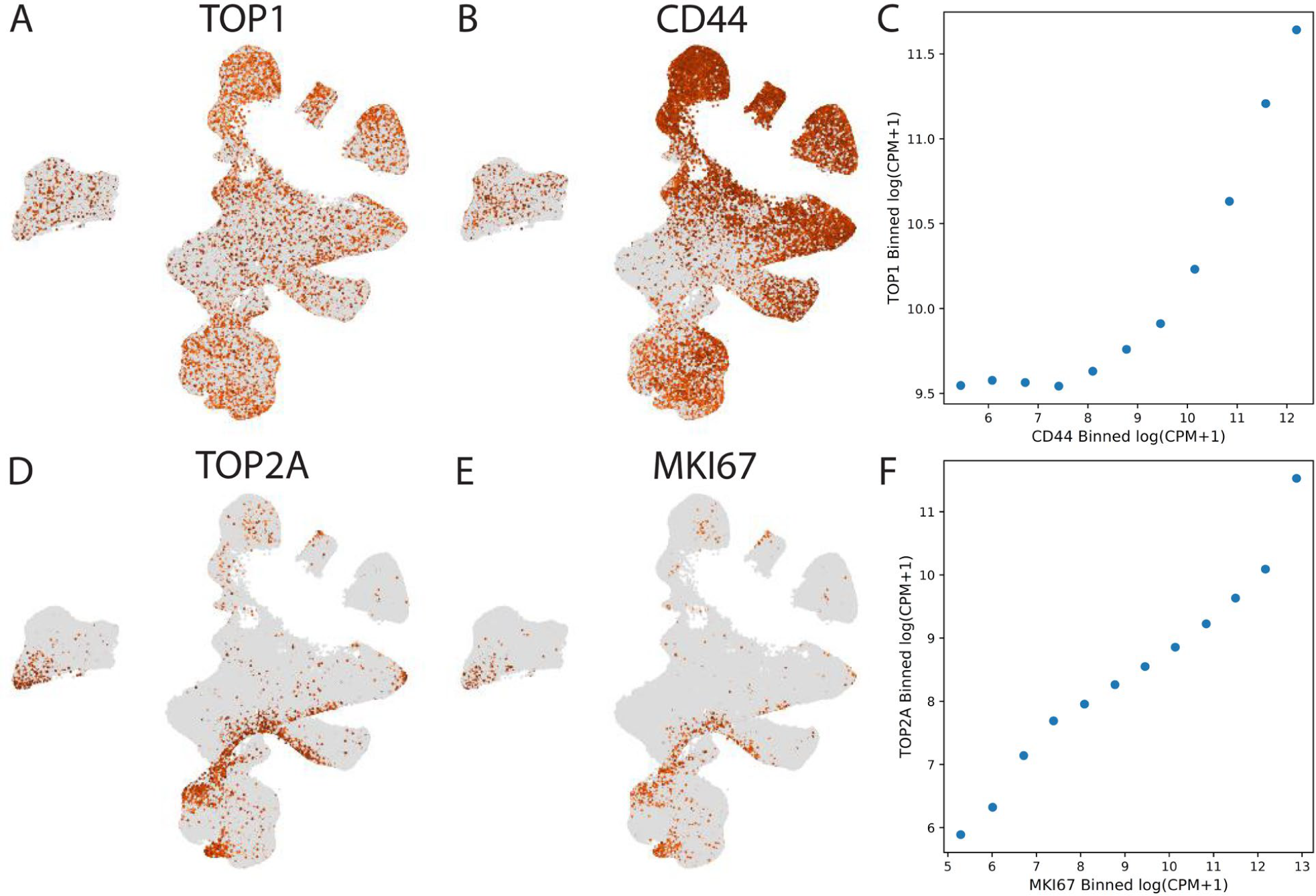
A) UMAP embeddings of the cell score matrix for the consensus scHPF model of transformed glioma cells colored by log(CPM+1) expression of *TOP1*; B) same as A) but for *CD44*; C) correlation between binned expression of *TOP1* and *CD44*; D) same as A) but for *TOP2A*; E) same as A) but for *MKI67*; F) same as C) but for *TOP2A* and *MKI67*.

#### Identification of Cell Type-Specific Drug Responses in the Glioma Microenvironment with Consensus scHPF

A critical advantage of *ex vivo* slice cultures is that they preserve the tumor microenvironment, allowing for investigation of cell type-specific drug responses in non-neoplastic brain and infiltrating immune cells. Myeloid cells are the most abundant infiltrating immune cell population in gliomas and can include both brain-resident microglia and bone marrow-derived macrophages. Microglia tend to exhibit a more pro-inflammatory phenotype in gliomas, whereas macrophages are thought to be more immunosuppressive and are associated with recurrence and poor survival^15,19^. **Figure 6** shows the effects of each drug on four key signatures derived from our myeloid-specific scHPF model. Similar to the transformed glioma cells, treatment with the HDAC inhibitors panobinostat and givinostat leads to significant upregulation of the highly inducible metallothionein gene cluster (**Figure 6A**). Consistent with our previous report, panobinostat treatment also strongly downregulates two factors enriched in macrophage-specific markers (**Figure 6B-C**)^8^. Thus, panobinostat may deplete or reprogram macrophages. Interestingly, while the other HDAC inhibitor in our study, givinostat, does not have this same effect, topotecan does. On the other hand, the effects of topotecan and panobinostat on a pro-inflammatory factor with genes that would typically be associated with microglia show no significant alterations across drugs (**Figure 6D**). These findings raise the possibility that topotecan and panobinostat might be effective at reprogramming the immune microenvironment of gliomas to a less immunosuppressive state.

**Figure 6.**
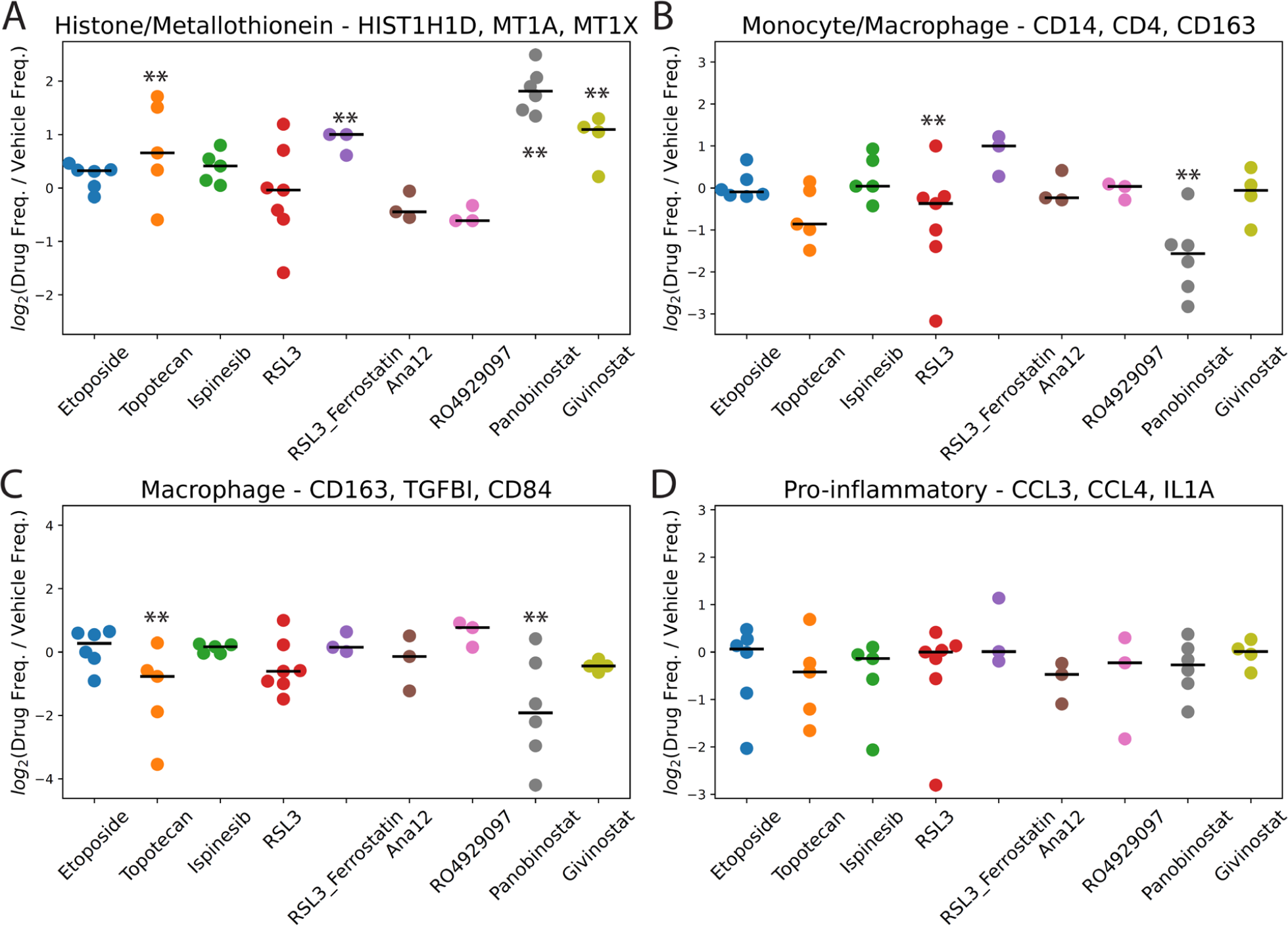
Fold-changes in the frequencies of cells with high cell scores in drug-treated vs. vehicle control-treated slice cultures in the myeloid cell scHPF model for A) a histone/metallothionein factor; B) a monocyte/macrophage factor; C) a macrophage factor; and D) a pro-inflammatory factor. Here, each dot represents an individual patient (i.e. biological replicates). For each drug, ** indicates FDR<0.05 based on a linear mixed model.

Overall, we found that the nine drugs tested here have more moderated and less consistent effects on oligodendrocytes, possibly because they are designed to target transformed cells. Nonetheless, we did observe two interesting effects on oligodendrocytes from the two HDAC inhibitors panobinostat and givinostat. Unlike in the transformed glioma cells and myeloid cells, where we observed metallothionein induction by both HDAC inhibitors, this effect appears restricted to panobinostat in oligodendrocytes (**Figure 7A**). More interestingly, we observed a consistent and significant upregulation of a factor marked by several sterol biosynthesis-related genes (*FAXDC2, DHCR7, TM7SF2, MVK*) by both HDAC inhibitors across all patients tested (**Figure 7B**). Cholesterol biosynthesis plays a key role in myelination by oligodendrocytes, with several of genes and the pathway in general showing strong upregulation in other neurological disorders such as multiple sclerosis, where remyelination is occurring^20^.

**Figure 7.**
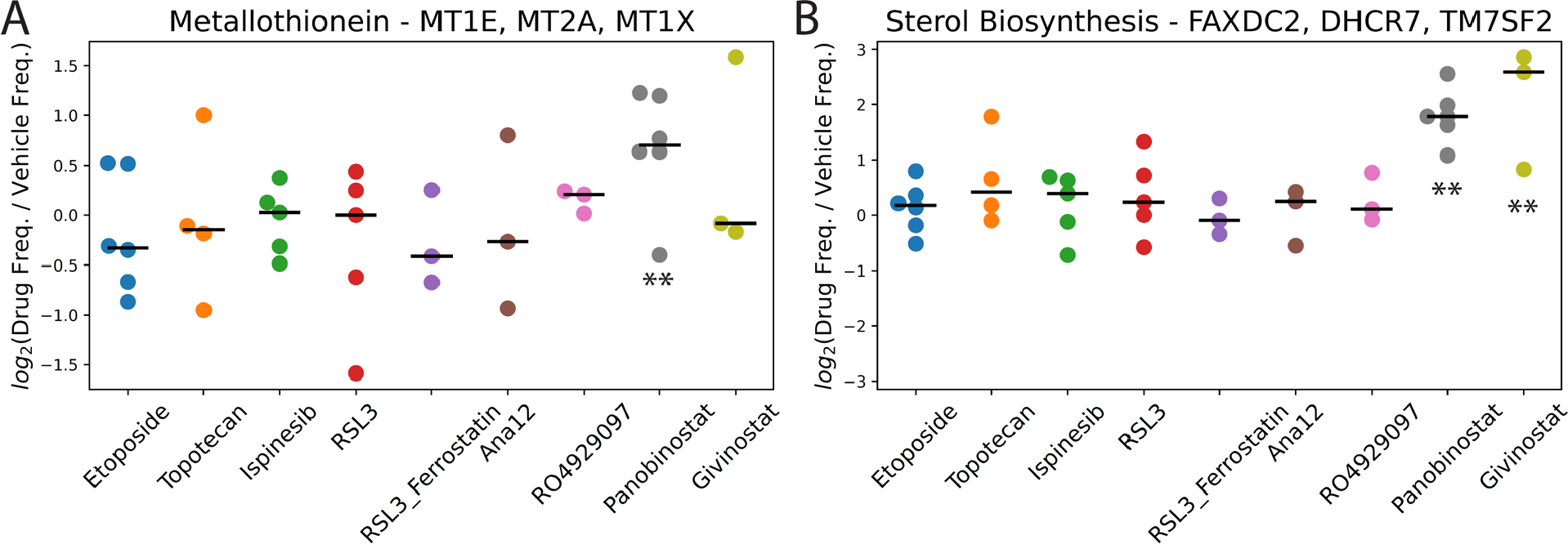
Fold-changes in the frequencies of cells with high cell scores in drug-treated vs. vehicle control-treated slice cultures in the oligodendrocyte scHPF model for A) a metallothionein factor and B) a sterol biosynthesis factor. For each drug, ** indicates FDR<0.05 based on a linear mixed model.

## Discussion

Consensus scHPF produces probabilistic factor models that integrate complex scRNA-seq datasets including multiple individuals, experimental conditions, and replicates. By identifying factors that occur reproducibly across models, consensus scHPF provides a principled approach to determining an appropriate value of *K*, the number of factors in the final, consensus model. This results in robust and interpretable factors that describe major gene co-expression patterns across a population of cells.

In developing consensus scHPF, we identified a number of measures that can improve performance in some cases. For example, for datasets with highly disparate cell numbers or coverage across conditions, balancing the cell numbers and counts per cell across the relevant covariate by random sub-sampling can be beneficial. The consensus scHPF release (https://github.com/simslab/consensus_scHPF_wrapper) provides scripts to assist users with these tasks along with a wrapper that parallelizes multi-model generation using the original scHPF software and performs consensus clustering and model refinement.

We applied consensus scHPF to a large scRNA-seq dataset comprised of ∼400,000 cells profiled from *ex vivo* slice cultures of human glioma surgical specimens across 19 patients and 10 drug perturbations (for a total of 52 unique samples). Because differences in cellular linages of these complex tumors are a dominant source of gene expression variance, we constructed consensus scHPF models separately for three major cellular populations – the transformed glioma cells, myeloid cells, and oligodendrocytes. As described in detail above, these models corroborated results from two of our previous, more focused studies^8,9^ and revealed some surprising findings, particularly with respect to the cell type-specific effects of topotecan. We found that topotecan behaves very differently from etoposide, the other topoisomerase poison in our dataset, in both transformed glioma cells and myeloid cells. We further found that the expression patterns of the genes encoding their respective targets, namely *TOP1* and *TOP2A*, are dramatically different and may explain the observed discrepancy. These findings are particularly timely, because of the recently completed clinical trial testing local delivery of topotecan in GBM by chronic convection enhanced delivery^21^. RNA-seq analysis of the pre- and post-treatment specimens from this trial suggested that long-term treatment with topotecan can also deplete proliferating cells. Thus, chronic local delivery of topotecan may target a broad spectrum of cellular states in GBM, including both transformed glioma cells and macrophages.

Overall, we anticipate that consensus scHPF will see widespread use for *de novo* gene signature identification from complex, integrated scRNA-seq datasets. These models will greatly aid in interpretation, particularly when the data are not well-described by clustering or contain multiple cell state transitions. Finally, we hope that our large-scale drug perturbation dataset from experiments in complex human surgical tissues will provide insights into cell type-specific drug responses beyond what we describe here and serve as a valuable resource for further methods development.

## Methods

### Clustering Factors in Consensus scHPF

As described above, we run scHPF with multiple (typically 5-10) random initializations for multiple values of *K* and select the top *m* models for each value of *K* with the lowest mean negative log likelihood on the training data. For each of the *N*_*m*_ models that resulted, we calculated each factors’ gene scores, and concatenated all factors (columns) across all models into a matrix *X* with factors as columns, genes as rows, and values set to scHPF gene scores. We then reduced this to a submatrix *X*_*CV*_, which contained the 1,000 rows (genes) with the highest coefficient-of-variation (CV). We note that the 1,000 genes with the highest CV did not correspond to genes with the highest mean due to the built-in normalization in scHPF’s gene scores.

To identify factors that were reproduced across multiple models, we clustered the factors using the Walktrap algorithm on a k-nearest neighbors graph of the columns of *X*_*CV*_. Reasoning that the patterns should be reproduced across at least a quarter of randomly initialized models, we set the number of neighbors to be *int(0*.*25N*_*m*_), and constructed a k-nearest neighbors graph using Pearson’s correlation distance. We ran the Walktrap algorithm on this graph using the *community_walktrap* function (python-igraph v0.8.2) with weights equal to the Jaccard similarity between adjacent nodes’ neighbors and default parameters otherwise. We selected the number of clusters by examining the partitions’ modularity (calculated using *igraph*.*clustering*.*VertexDendrogram*.*modularity*) as a function of the number of clusters, and set the number of partitions to the center of the peak in modularity.

### Procurement of Human Glioma Surgical Specimens

All tumor specimens collected were de-identified and under the approval of the Columbia University Irving Medical Center Institutional Review Board. Clinical metadata can be found in **Table S1**.

### Ex vivo Slice Culture and Drug Perturbation in Human Glioma Surgical Specimens

Collected tumor specimens were prepared and processed for *ex vivo* slice culture followed by drug perturbation as described previously (ref Zhao *et al*. 2021). Briefly, tumor specimens were kept in ice-cold artificial cerebrospinal fluid (ACSF) solution immediately after surgical removal and sliced using a tissue chopper (McIlwain) at a thickness of 500 μm under sterile conditions. Generated slices were first transferred to 6-well plates and kept in ice-cold ACSF solution followed by a 15 minute recovery to reach room temperature. Then we placed each slice on top of a porous membrane insert (0.4 μm, Millipore) sitting in 6-well plates and added 1.5 mL maintenance medium consisting of F12/DMEM (Gibco) supplemented with N-2 Supplement (Gibco) and 1% antibiotic-antimycotic (ThermoFisher) to the bottom of each well and 10μl maintenance medium directly on top of each slice. After culturing slices in a humidified incubator at 37°C and 5% CO_2_ for 6hr, we replaced the medium with pre-warmed medium containing drugs with desired concentration (listed in **Table S2**) or corresponding volume of vehicle (DMSO) then cultured slices in a humidified incubator at 37°C and 5% CO_2_ for 18 hrs.

### Slice Culture Dissociation

At the end of drug treatment, tissue slices were dissociated into single cell suspensions for microwell-based single-cell RNA-seq^22^. Tissue slices derived from TB5884, TB5886, TB5944, TB5966, TB5974, TB5980, TB6140, TB6181, TB6224, TB6328, TB6393, and TB6458 were dissociated as described in Zhao *et al*. using the Adult Brain Dissociation kit (Miltenyi Biotec) on gentleMACS Octo Dissociator with Heaters (Miltenyi Biotec) according to the manufacturer’s instructions. Tissue slices derived from TB6528, TB6534, TB6545 (1 RSL3 and 1 vehicle slice), TB6813, and TB6814 were dissociated as described in Banu *et al*.^*9*^ using the Papain Dissociation System. TB6488, TB6505 and TB6545 (1 givinostat and 7 vehicle slices) were dissociated using the Adult Brain Dissociation kit (Miltenyi Biotec) with modifications. Briefly, dissociation buffer was prepared freshly according to the manufacturer’s instructions. Each tissue slice was collected into a well of a 12-well plate and washed in ice-cold Dulbecco’s phosphate-buffered saline (D-PBS) with calcium, magnesium, glucose, and pyruvate (Lonza) following a 30 min incubation in 1ml of dissociation buffer at 37°C in a shaking incubator at 600rpm. Dissociated cells of each slice were collected with 6ml cold D-PBS and apply to a MACS SmartStrainer (70 μm, Miltenyi Biotec) placed on one well of a deepwell 24-well plate following centrifugation at 300×g for 10 minutes at 4°C. The cell pellet was then processed for debris removal and red blood cell removal according to the manufacturer’s instructions.

### scRNA-seq in Microwell Platform

Following dissociation, we used our previously described microwell-based platform to perform scRNA-seq from each slice culture^23,24^. During library construction, pooled cDNA libraries from each slice culture were associated with a unique Illumina index sequence to facilitate pooled sequencing of all of the slice cultures from a given patient. The resulting pooled libraries were either sequenced an Illumina NextSeq500/550 (8-cycle index read, 26-cycle read 1 containing the cell barcode (CB) and unique molecular identifier (UMI), 58-cycle read 2 containing the transcript sequence) or an Illumina NovaSeq 6000 (8-cycle index read, 26-cycle read 1, 151-cycle read 2).

### scRNA-seq Data Processing

scRNA-seq data were processed as described previously^8^. Briefly, after trimming and aligning the raw reads, we corrected any datasets obtained using the Illumina NovaSeq 6000 for index swapping using the algorithm of Griffiths et al^25^. We then assigned read addresses compressed of a CB, UMI, and aligned gene to each read, collapsed reads with duplicate addresses, and corrected errors in the CB and UMI. Finally, we identified CBs that were likely to originate from cells using the EmptyDrops algorithm^26^ and applied several quality control filters to the resulting CBs as described in Zhao et al^8^ to arrive at a final count matrix for each sample.

### Identification of Major Cell Types from scRNA-seq

Major cell types were identified as described previously^8^. Briefly, we first merged scRNA-seq data of all samples derived from the same patient for unsupervised clustering analysis (www.github.com/simslab/cluster_diffex2018)^5^. We used Louvain community detection as implemented in Phenograph for unsupervised clustering with k=20 for all k-nearest neighbor graphs^27^. The marker genes was identified using the drop-out curve method as described in previously^5^ for each individual sample and took the union of the resulting marker sets to cluster and embed the merged dataset. We defined putative malignant cells and non-malignant cells using the genes most specific to each cluster. Putative tumor-myeloid doublet clusters were removed prior to malignant analysis. Next, we computed the average gene expression on each somatic chromosome as described in Yuan *et al*^*15*^. For data obtained from glioblastoma tissues (IDH1 wild type tissues), we define the malignancy score to be the log-ratio of the average expression of Chr. 7 genes to that of Chr. 10 genes as described previously. For data obtained from the TB6505, Chr. 2 amplification and Chr. 13 deletion were observed from the whole-genome sequencing results, therefore we define the malignancy score to be the log-ratio of the average expression of Chr. 2 genes to that of Chr. 13 genes. For data obtained from the TB6505, we define the malignancy score to be the log-ratio of the average expression of Chr. 2 genes to that of Chr. 1 genes as described in Banu *et al*.^*9*^. We plotted the distribution of malignancy score and fit a two-component Gaussian mixture model to the malignancy score distribution and established a threshold at 1.96 standard deviations below the mean of the Gaussian with the higher mean (i.e. 95% confidence interval). Putative malignant cells with malignancy scores below this threshold and putative non-tumor cells with malignancy scores above this threshold were discarded as non-malignant or potential multiplets. The malignant analysis of newly reported tissues was shown in **Figure S1**. We further manually annotated non-malignant cells into myeloid cells (*CD14, AIF1, TYROBP, C1QA*), oligodendrocytes (*PLP1, MBP, MAG, SOX10*), T cells (*TRAC, TRBC1, TRBC2, CD3D*), endothelial cells (*ESM1, ITM2A, CLDN5*), and pericytes (*PDGFRB, COL3A1, RGS5*) based on the highly enriched marker genes of each cluster.

### Consensus scHPF Models of Drug-Perturbed Human Glioma Slice Cultures

For each major cell type (transformed glioma cells, myeloid cells, and oligodendrocytes), we constructed a merged count matrix in loom format using *loompy*. Next, for each cell type, we used the *get_training_test_looms*.*py* script to generating a loom file for the cells from which the consensus scHPF model would be trained and a smaller loom file for cells comprising the test set with 50 cells per patient. We then reformatted the training and test set loom files to the appropriate sparse matrix format for running scHPF using the *scHPF prep* and *scHPF pre-like* commands, respectively. For *scHPF prep*, we used a white list comprising protein-coding genes and excluding T cell and immunoglobulin receptor genes. We required genes to be detected in 1% cells to be included in the model. Finally, we used the *scHPF_consensus*.*py* script to run consensus scHPF. We generated scHPF models for each value of *K* from 15-30 with five trials (parameter *n*) per value of *K*. For Walktrap clustering, we required clusters to contain factors from at least two models (parameter *m*). This resulted in consensus scHPF models containing 22, 24, and 17 factors for transformed glioma cells, myeloid cells, and oligodendrocytes, respectively. We used the *scHPF score* command to compute cell and gene score matrices for each factor in each of the three consensus models.

### Analysis of Consensus scHPF Models to Identify Cell Type-Specific Drug Responses

For each patient-drug sample, we compared the cell scores for each factor between the scRNA-seq data for the drug-treated slice and the corresponding vehicle control-treated (DMSO) slice(s) from the same patient. To make this comparison, we computed the median absolute deviation (MAD) cell score for each factor and calculated the frequency of cells in each sample that were more than two MADs above the median cell score for the factor. The values plotted in **Figures 4, 6, 7** are the log-scaled fold-changes for this cell frequency between the drug-treated slice and vehicle control-treated slice(s) for a given patient. For each drug-factor combination, we calculated FDR-corrected p-values using a linear mixed model with the log-scaled fold-change in cell frequency as the response variable and the drug treatment as a categorical covariate. We treated patients as random effects. Coefficients and p-values for each covariate were computed using the *MixedLM* function in the Python package *statsmodels*. P-values were corrected for false discovery using the *multipletests* function in *statsmodels* with the Benjamini-Hochberg procedure.

## Data and Code Accessibility

The raw count matrix and metadata for the entire integrated dataset used in these studies is available as a loom file at:

https://drive.google.com/file/d/18-KInmm43wKdBX95Gq9xbuzAQwtLjgE9/view?usp=sharing

The original source code with tutorials for scHPF, which is used to build individual models in consensus scHPF can be found at https://github.com/simslab/scHPF. Code for running consensus scHPF along with helper scripts and instructions can be found at https://github.com/simslab/consensus_scHPF_wrapper.

## Acknowledgements

P.A.S. was supported by the Mark Foundation for Cancer Research Grant MFCR18-019 ELA. P.A.S., J.N.B., and P.C. were supported by NIH/NINDS R01NS103473. This research was funded in part through the NIH/NCI Cancer Center Support Grant P30CA013696, which supports the Genomics and High Throughput Screening Shared Resource. We thank Dr. Peter Szabo for creating Figure 1.

