## Supplementary Figures and Tables for "Consensus scHPF Identifies Cell Type-Specific Drug Responses in Glioma by Integrating Large-Scale scRNA-seq"

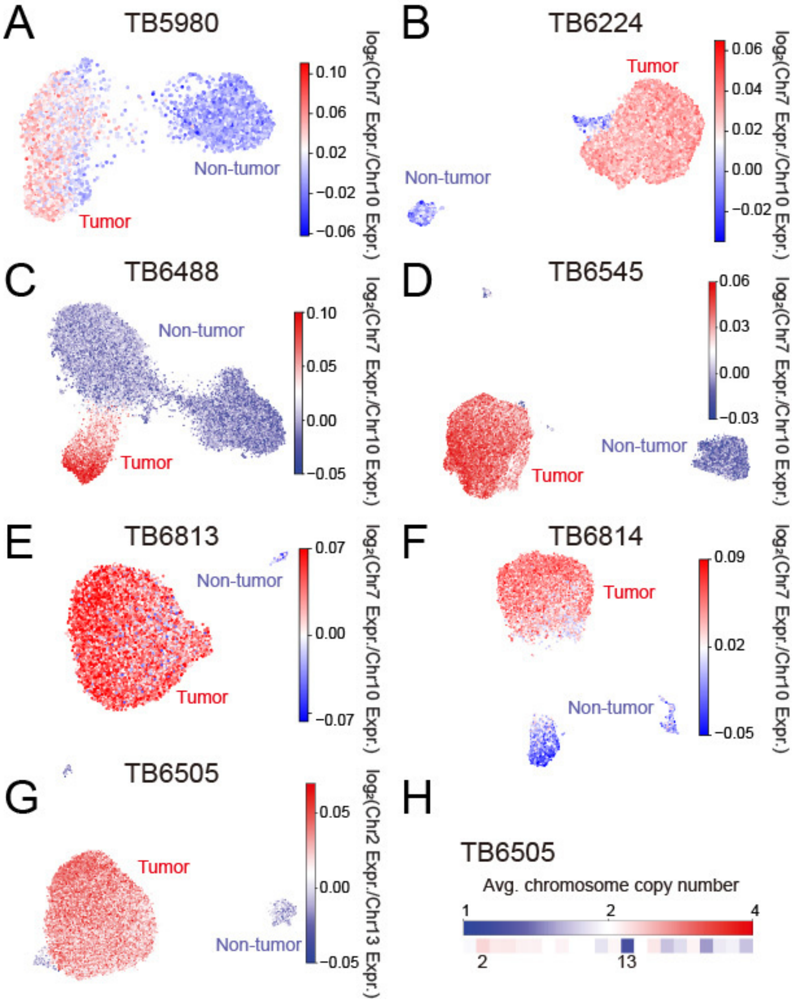


**Figure S1.** A)-F) UMAP embeddings of scRNA-seq data from GBM patient slice cultures that were not reported in previous publications colored by the log-ratio of the average expression of genes on Chr. 7 to Chr. 10. Chrs. 7 and 10 are pervasively amplified and deleted, respectively, in GBM, and so their expression ratio can be used to identify clusters that correspond to transformed glioma cells. G) UMAP embedding of scRNA-seq data from slice cultures from an IDH1mt Grade IV astrocytoma, which does not harbor amplification of Chr. 7 and loss of Chr. 10. Based on low-pass WGS of the corresponding tumor tissue, which identified amplification of Chr. 2 and loss of Chr. 13, as shown in the heatmap in H), we used the expression ratio of these two chromosomes to identify transformed glioma cells in this patient.

### Supplementary Tables

**Table S1.** Summary of patients, specimens, and culture conditions.

| Tissue ID | Age Range | Sex | Location | Diagnosis | IDH1 Status | EGFR status | Samples | Reference |
|---|---|---|---|---|---|---|---|---|
| TB6488 | 50-60 | F | Left temporal | Glioblastoma, WHO Grade IV | Wildtype | Amplified | 7 vehicle slices, 1 givinostat slice | This study |
| TB6140 | 60-70 | M | Right temporal | Glioblastoma, WHO grade IV | Wildtype | Amplified | 6 vehicle slices, 1 panobinostat slice | PW040, Zhao *et al.*, 2021 |
| TB5884 | 60-70 | M | Right parietal | Glioblastoma, WHO grade IV | Wildtype | Unamplified | 2 vehicle slices, 1 etoposide, 1 panobinostat, 1 Ana12, 1 ispinesib, and 1 RO492997 slice | PW030, Zhao *et al.*, 2021 |
| TB5966 | 60-70 | F | Left parieto-occipital | Glioblastoma, WHO grade IV | Wildtype | Unamplified | 2 vehicle slices, 1 etoposide and 1 panobinostat slice | PW034, Zhao *et al.*, 2021 |
| | | | | | | | 1 Topotecan, 1 ispinesib, and 1 RO4929097 slice | This study |
| TB6505 | 30-40 | M | Left frontal | Astrocytoma, WHO Grade IV | Mutant | Unamplified | 7 vehicle slice, 1 givinostat slice | This study |
| TB6528 | 60-70 | F | Left temporal | Glioblastoma, WHO Grade IV | Wildtype | Unamplified | 1 vehicle slice, 1 RSL3 slice | Banu *et al.*, 2023 |
| TB5944 | 60-70 | M | Left frontal | Glioblastoma, WHO grade IV | Wildtype | Amplified | 3 vehicle slices, 1 etoposide and 1 panobinostat slice | PW032, Zhao *et al.*, 2021 |
| | | | | | | | 1 Topotecan, 1 Ana12 and 1 ispinesib slice | This study |
| TB5974 | 50-60 | M | Right temporal | Glioblastoma, WHO grade IV | Wildtype | Amplified | 2 vehicle slices, 1 etoposide and 1 panobinostat slice | PW036, Zhao *et al.*, 2021 |
| | | | | | | | 1 Topotecan, 1 RO4929097, 1 ispinesib, and 1 Ana12 slice | This study |

| | | | | | | | | |
|---|---|---|---|---|---|---|---|---|
| TB6328 | 70-80 | F | Left frontal | Glioblastoma, WHO Grade IV | Wildtype | Amplified | 2 vehicle slices, 2 RSL3 slices, and 2 RSL3+Ferrostatine slices | Banu *et al.*, 2023 |
| TB6458 | 50-60 | M | Right frontal | Glioblastoma, WHO Grade IV | Wildtype | Amplified | 2 vehicle slices, 1 RSL3 slice, and 1 RSL3+Ferrostatine slice | Banu *et al.*, 2023 |
| TB6813 | 70-80 | F | Left parietal | Glioblastoma, WHO Grade IV | Wildtype | Amplified | 1 vehicle slice, 1 RSL3 slice | This study |
| TB5886 | 50-60 | F | Splenial glioma extension into left parietal | Glioblastoma, WHO grade IV | Wildtype | Amplified | 1 vehicle slice, 1 etoposide slice | PW029, Zhao *et al.*, 2021 |
| | | | | | | | 1 ispinesib slice | This study |
| TB6545 | 30-40 | M | Right frontal | Glioblastoma, WHO Grade IV | Wildtype | Unamplified | 1 vehicle slice, 1 RSL3 slice | Banu *et al.*, 2023 |
| | | | | | | | 7 vehicle slices, 1 givinostat slice | This study |
| TB6224 | 60-70 | M | Right posterior frontal lobe | Glioblastoma, WHO Grade IV | Wildtype | Amplified | 7 vehicle slices, 1 givinostat slice | This study |
| TB6814 | 60-70 | M | Right temporal | Glioblastoma, WHO Grade IV | Wildtype | Amplified | 1 vehicle slice, 1 RSL3 slice, 1 Topotecan | This study |
| TB6534 | 50-60 | F | Right temporal | Glioblastoma, WHO Grade IV | Wildtype | Unamplified | 1 vehicle slice, 1 RSL3 slice | Banu *et al.*, 2023 |
| TB5980 | 60-70 | M | Right medial temporal | Glioblastoma, WHO Grade IV | Wildtype | Amplified | 1 vehicle slice, 1 Topotecan slice | This study |
| TB6181 | 30-40 | F | Frontal and temporal | Anaplastic Astrocytoma, WHO Grade III | Mutant | Unamplified | 2 vehicle slices, 1 RSL3 slice, 1 RSL3+Ferrostatine slices | Banu *et al.*, 2023 |
| TB6393 | 70-80 | F | Right frontal | Glioblastoma, WHO grade IV | Wildtype | Unamplified | 4 vehicle slices, 3 etoposide slices, 3 panobinostat slices | Zhao *et al.*, 2021 |

**Table S2.** Summary of drugs and concentrations (including newly and previously reported studies).

| **Drug Name** | **Resource** | **Working Concentration** | **Reference** |
|---|---|---|---|
| Etoposide | Tocris Bioscience (Cat# 1226/100) | 2.5 µM | Zhao *et al.*, 2021 |
| Panobinostat (LBH589) | Selleck Chem (Cat # S1030) | 0.2 µM | Zhao *et al.*, 2021 |
| Ana12 | TOCRIS (Cat #4781 - 10 mg) | 40 nM | Zhao *et al.*, 2021 |
| Ispinesib | Selleck Chem (Cat# S1452) | 1.8 nM | Zhao *et al.*, 2021 |
| RO492997 | Selleck Chem (Cat# S1575) | 50 nM | Zhao *et al.*, 2021 |
| RSL3 | Brent Stockwell Lab | 50 nM | Banu *et al.*, 2023 |
| Ferrostatin | Brent Stockwell Lab | 10 µM | Banu *et al.*, 2023 |
| Givinostat | Sigma-Aldrich (Cat# SML1772-25MG) | 0.23 µM | This study |
| Topotecan | Selleckchem (Cat# S1231) | 20 µM | This study |
